# The role of evolutionary modes for trait-based cascades in mutualistic networks

**DOI:** 10.1101/2021.10.05.462726

**Authors:** Vinicius Augusto Galvão Bastazini, Vanderlei Debastiani, Laura Cappelatti, Paulo Guimarães, Valério D. Pillar

**Affiliations:** Rui Nabeiro Biodiversity Chair, MED - Mediterranean Institute for Agriculture, Environment and Development, University of Évora. Casa Cordovil 2° Andar, Rua Dr. Joaquim Henrique da Fonseca, 7000 – 890 Évora, Portugal; (Graduate Program in Ecology, Universidade Federal do Rio Grande do Sul, Porto Alegre, RS, 91501-970, Brazil); Graduate Program in Ecology, Universidade Federal do Rio Grande do Sul, Porto Alegre, RS, 91501-970, Brazil; Independent researcher, Helsinki, Finland; Departmento de Ecologia, Instituto de Biociências, Universidade de São Paulo, Rua do Matão, Travessa 14, no. 321, 05508-900, São Paulo, SP, Brazil; Department of Ecology, Universidade Federal do Rio Grande do Sul, Porto Alegre, RS, 91501-970, Brazil

## Abstract

The erosion of functional diversity may foster the collapse of ecological systems. Functional diversity is ultimately determined by the distribution of species traits. As species traits are a legacy of species evolutionary history, one might expect that the mode of trait evolution influences community resistance under the loss of functional diversity. In this paper, we investigate the role of trait evolutionary dynamics on the robustness of mutualistic networks undergoing the following scenarios of species loss: i) random extinctions, ii) loss of functional distinctiveness and iii) extinctions biased towards larger sizes. We simulated networks defined by models of single trait complementary and evolutionary modes where traits can arise in recent diversification events with weak phylogenetic signal, in early diversification events with strong phylogenetic signal, or as a random walk through evolutionary time. Our simulations show that mutualistic networks are especially vulnerable to extinctions based on trait distinctiveness and more robust to random extinction dynamics. The networks show an intermediate level of robustness against size-based extinctions. Despite the small range of variation in network robustness, our results show that the mode of trait evolution matters for network robustness in all three scenarios. Networks with low phylogenetic signal are more robust than networks with high phylogenetic signal across all scenarios. As a consequence, our results predict that mutualistic networks based upon current adaptations are more likely to cope with extinction dynamics than those networks that are based upon conserved traits.

## Introduction

Understanding how ecological systems respond to disturbances is a central and long-standing issue in theoretical and applied ecology (May 1972, 2001; Pimm 1984; Neutel et al. 2002; Allesina & Tang 2012; Myers et al. 2015; Pires et al. 2015). Given the pace of anthropogenic-induced mass extinction, with species dying out three orders of magnitude faster than the extinction background rate inferred from fossil record (Pimm et al. 2014; Ceballos et al. 2015), the need to understand and predict species extinctions has become a fundamental task to mitigate human impact on ecosystems (Vieira & Almeida-Neto 2014; Ceballos et al. 2015). Ecologists have long acknowledged that the species loss may trigger cascading effects in ecological communities, which might bring other species to extinction and even entire ecosystems to collapse (Estes et al. 1998; Jackson et al. 2001; Colwell et al. 2012; Säterberg et al, 2013; Vieira et al. 2013; Brodie et al. 2014; Vieira & Almeida-Neto 2014). However, to date, most studies that have examined the magnitude of biodiversity loss usually ignore co-extinction processes (Dunn et al. 2009; Vieira & Almeida-Neto 2014; but see Silva et al. 2007 and Strona & Bradshaw 2018).

Mutualistic networks are formed by sets of interacting species, generating mutual benefits for participant species (Bronstein 2001; Bascompte & Jordano 2007, 2014). Mutualistic networks include a wide range of taxonomic groups and interaction types, such as interactions between flowering plants and their animal pollinators and seed dispersers (e.g., Bascompte & Jordano 2007; Muller-Landau & Hardesty 2005; Vizentin-Bugoni et al. 2014), animal cleaning associations (e.g., Wicksten 1998; Guimarães et al. 2007; Sazima et al. 2010) and many forms of human-microbe interactions (Dethlefsen et al. 2007). Mutualistic interactions provide an important model system for understanding properties of ecological communities given their paramount role in shaping ecoevolutionary dynamics, biodiversity patterns, ecosystem functioning (Ferriere & Legendre 2013; Bascompte & Jordano 2014; Scheuning et al. 2015, Guimarães Jr 2020) and, consequently, for their importance to the development of conservation strategies (Kiers et al. 2010; Brodie et al. 2014). Although mutualists can be flexible with regards to their partners (Bascompte and Jordano 2014), extinctions in mutualistic systems have the potential to accelerate biodiversity loss and ecosystem disruption (Kiers et al. 2010).

Among the factors that are recognized as important drivers of mutualistic network organization, species functional traits play a crucial role. Functional traits are behavioral, morphological or ecological characteristics associated with organismal fitness, biotic interactions and/or an ecosystem function of interest (Schmitz et al. 2015; Lefcheck et al. 2015). Functional traits are critical to network organization because they can directly constrain or enable the likelihood of an interaction among two or more individuals, imposing thresholds on trait values for feasible interactions (Santamaría & Rodríguez-Gironés 2007; Vizentin-Bugoni et al. 2014; Minoarivelo & Hui 2016; Bastazini et al. 2017, Guimarães Jr 2020).

Species traits may also affect extinction probability, as taxa with some specific traits, such as large body size, and narrow niche breadth are especially more prone to extinction (Purvis et al. 2000; Cardillo et al. 2005; Reynolds et al. 2005; but see Chichorro et al. 2019). The robustness of ecological networks, i.e., the system’s tolerance to species loss, has been traditionally evaluated based on scenarios where secondary extinctions are driven by species specialization (i.e., number of interacting partners) and/or on stochastic processes (Solé & Montoya 2001; Dunne et al. 2002; Memmot et al. 2004; Burgos et al. 2007; Rezende et al. 2007a; Pocock et al 2012 but see Vidal et al. 2013). These studies help us to broaden our understanding of ecological resistance. A next step in the analysis of network vulnerability is to explore the role of how ecological and evolutionary factors affect the likelihood of species becoming extinct (Bastazini et al. 2019; but see Vieira et al. 2013; Astegiano et al. 2015). For example, as trait redundancy may play an important role in network robustness, extinctions are expected to have a small effect on robustness if all species are functionally similar (higher trait resemblance), but a large effect if species have different trait values (Fonseca & Ganade, 2001).

As species traits are largely a legacy of their evolutionary history (Grafen 1989; Diniz-Filho et al. 2012; Mouquet et al. 2012), it is expected that the mode of evolution, i.e., how traits arise along the phylogenetic history of a clade (Burin et al. 2021), may play a pivotal role in ecological dynamics, and consequently, the robustness of networks that are losing functional diversity. Furthermore, recent evidence suggests that the loss of functional trait diversity takes a larger toll in ecological communities than taxonomic loss alone, making them more likely to collapse (Galetti et al. 2013; Brodie et al. 2014; Valiente-Banuet et al. 2015; Bastazini et al. 2019; Crooke et al. 2020).

Here, we theoretically explore how different modes of trait evolution may affect the robustness of mutualistic networks undergoing three distinct extinction scenarios of species loss: i) random extinctions, which serves as a baseline scenario; ii) loss of functional distinctiveness (i.e., species disappearing sequentially as a function of their functional distinctiveness); and iii) size trait (i.e. species with larger size-related traits disappearing first). As phylogenetically related species tend to interact with a similar set of species (Rezende et al. 2007a), we predicted that networks formed by species with higher levels of phylogenetic signal in traits would be more robust to secondary extinction, as a result of a more cohesive and redundant structure within the network.

## Modeling Approach and Statistical Analysis

### Eco-evolutionary Dynamics

We modeled the evolutionary dynamics of bipartite mutualistic networks, formed by two sets of interacting species (Fig. 1). We first produced simulated ultrametric phylogenetic trees of different sizes for each set of species, resulting from a uniform birth-death process (Nee et al. 1994). The size of simulated phylogenetic trees ranged from 10 to 20 species, which generated networks that varied in size, ranging from 20 to 40 species.

**Fig. 1.**
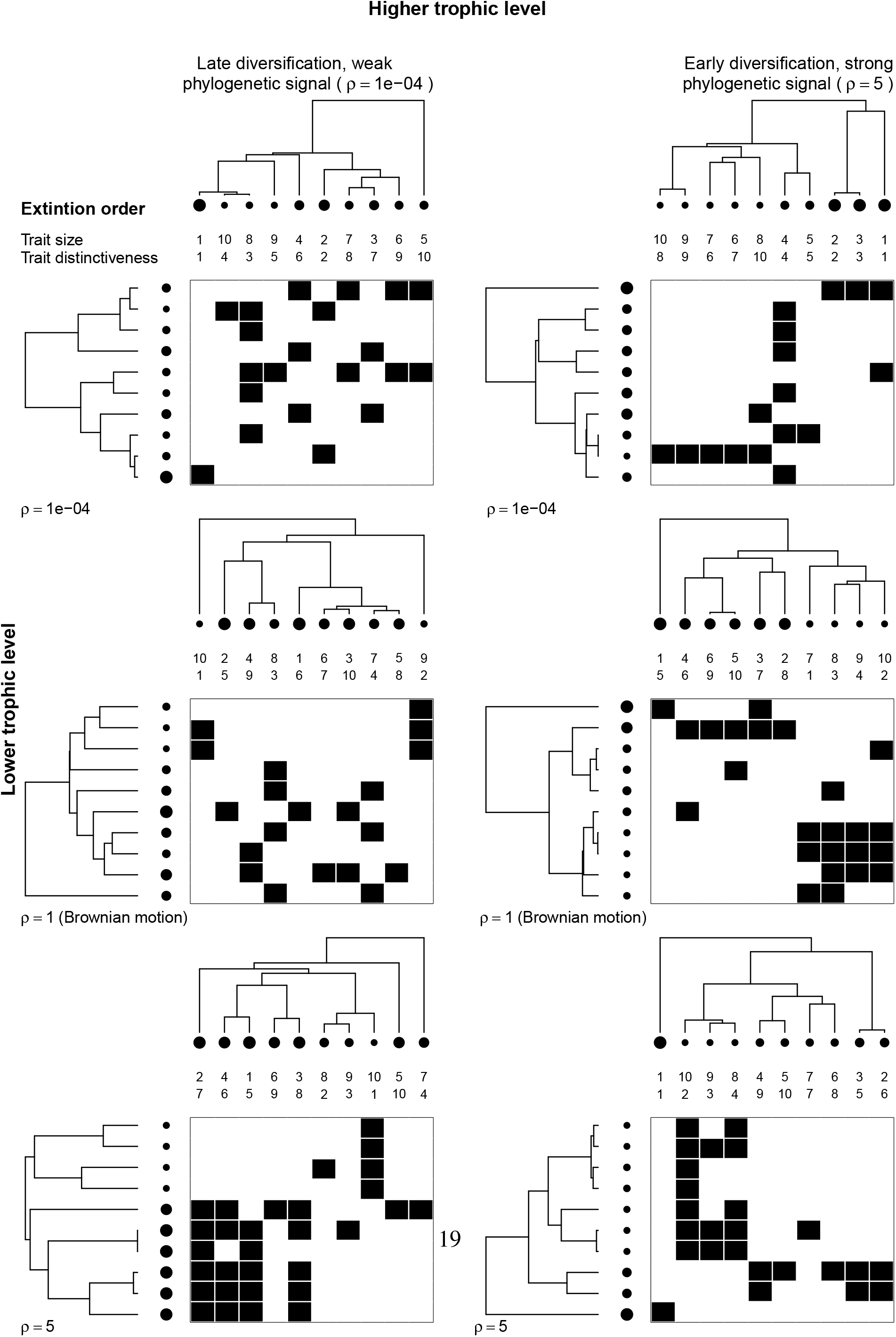
Examples of some of the possible eco-evolutionary dynamics of bipartite mutualistic networks we adopted in our simulations (six out of the 16 possible combinations). Species interactions are denoted in the bi-adjacency matrices along with their evolutionary trees and traits (black circles). The size of each circle corresponds to trait values. Graphen’s ρdefine the tempo and mode of trait evolution. A single-trait complementarity model defines the probability of interaction of two species. We assigned a random interaction to species that did not have any overlapping trait, to ensure all species interacted with at least one from the other set of species. More details are in the main text.

Secondly, we simulated the evolution of a single trait using a family of power transformations to the branch lengths of simulated phylogenetic trees (Grafen 1989). These transformations were achieved by raising the height of each phylogenetic tree to a different power, denoted by ρ (Grafen 1989). The range of powers used in these transformations simulates different evolutionary models (Fig. 1). When the height of a phylogenetic tree is raised to the power of 1, it simulates trait evolution under Brownian motion, as if evolution of traits followed a random walk through evolutionary time (Diniz-Filho et al. 2012), which implies that traits divergence increases linearly with time. Power values smaller than 1 compress deeper branch lengths, and expand them near the tips of the tree, simulating a recent diversification of traits with low phylogenetic signal (Diniz-Filho et al. 2012), while ρ values larger than 1 increase branch lengths near the root of the tree and simulate early diversification of traits with high phylogenetic signal (Diniz-Filho et al. 2012).

Networks were then generated using the single-trait complementarity model, given by equations 1 and 2, proposed by Santamaría & Rodríguez-Gironés (2007) which assumes that interactions between species can be described by a single trait. This approach emulates ecological systems such as pollination networks formed by flowering plants and birds, in which species interactions can be predicted by flowers’ corola length and hummingbird’s bill length (Vizentin-Bugoni et al. 2014, 2020; see also: Garibaldi et al. 2015, Stang et al. 2009, Donoso et al. 2017). Following Santamaría & Rodríguez-Gironés (2007) approach, a mean trait value and its variability characterize each species in the network and a pair of species is more likely to interact if their trait values overlap. In their definition,*V*_*i*_ and *W*_*j*_ is the central trait value for species *i* in one set (e.g, flowering plants) and species *j* in the other set (e.g., pollinators), respectively, and *δV*_*i*_ and *δW*_*j*_ are the range of variability of each trait for species i and j. Then, the value of each cell in the bi-adjacency matrix, corresponding to this pair of species *I*_*ij*_ will be

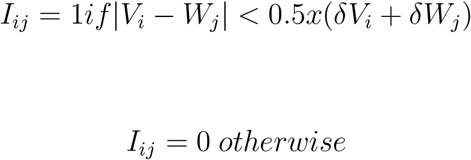

which means that a pair of species interact if the cell value is equal to one and they do not interact if it is equal to zero. The variability represented by *δV*_*i*_ and *δW*_*j*_were defined as random variables with uniform distributions in the intervals 0 – 0.25 (Santamaría & Rodríguez-Gironés 2007). To ensure all species within the network interacted at least with one species from the other set of species, we assigned a random interaction to species that did not have any overlapping trait (Fig. 1).

### Co-extinction analyses

We estimated network robustness (*R*) based on the area below the Attack Tolerance Curve (ATC; Albert & Barabási 2002; Memmott et al. 2004; Burgos et al. 2007). The ATC is a quantitative description of the network robustness measuring its ability to maintain its structural connectivity as species go extinct. The ATC is contained in the unit square and starts at a value 1 in the *y* -axis, when no species in one set of species are eliminated and all the species in the other set survive. As species are eliminated, the curve decreases monotonically to 1 in the *x* -axis as no species in one set survives because all the species in the other set went extinct (for further details see Burgos et al. 2007). *R* values closer to 1 indicate higher network robustness, i.e., the system is more tolerant to species extinctions. We used three distinct species elimination scenarios. First, we removed species based on their trait distinctiveness, which means that at each time step, the species with the most distinct trait value is eliminated (Bastazini et al. 2019). We estimated the functional distinctiveness of each species, following the approach proposed in Bastazini et al. (2019), using an analogous metric used in phylogenetic studies (Redding et al. 2008). Therefore, we built a functional dendrogram based on species trait resemblance and then calculated the functional distinctiveness of each species, defined as the sum of all edge lengths between the species and the root of the dendrogram, with each edge length divided by the number of species in the cluster it subtends (Bastazini et al. 2019). In the second elimination scenario, we simulated species extinctions based on size trait values, eliminating species with larger sizes, as empirical evidence suggest that larger species have a higher chance of dying out (Cardillo et al. 2005, Reynolds et al. 2005; Donoso et al. 2017; but see Chichorro et al. 2019). At last, we eliminated species at random from the higher trophic level, which serves as a baseline scenario to compare the effects of the two functional extinction scenarios.

We compared four evolutionary modes under four distinct scenarios. The four models compared are related to the evolutionary mode generating traits in the species belonging to the higher trophic level (e.g., pollinators, seed dispersers): i) Traits of species diversified recently, presenting low phylogenetic signal (ρ = 1e-04); ii) Trait evolution follows a Brownian process, (ρ = 1.0); and two models, where traits diversified in the beginning of the evolutionary process, with strong phylogenetic signal (ρ = 2.0 and 5.0, modes iii and iv, respectively). These models were compared in four distinct scenarios according with the evolution of traits in species in the set of species belonging to the lower trophic level (e.g., flowering plants): A) a random combination of evolutionary modes, where phylogenetic signal varies from low (ρ = 1e-04) to high phylogenetic signal (ρ = 5.0); B) late diversification of traits (ρ = 1e-04); C) trait evolution following a Brownian process, (ρ = 1.0); and D) early diversification of traits (ρ = 5.0).

We compared the effects of the evolutionary modes, within each scenario of species extinction, using a Bayesian analysis of variance, based on Jeffreys non-informative priors (Kinas & Andrade 2010). Based on the posterior distribution we calculated 95% Bayesian Credible intervals for each scenario. The posterior distributions of parameters are defined as:

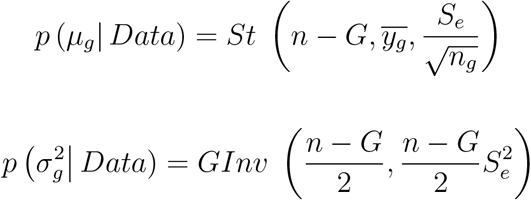

where *G* is the fixed factor representing the evolutionary modes, μ_*g*_ is the mean robustness response for each scenario, and σ^2^ is the variance (for further details see Kinas & Andrade 2010). The posterior distribution is simulated first sampling σ^2^_*g*_. Then, sampling values are taken from a multivariate normal distribution with μ_g_ equal to the mean robustness, with covariance matrix giving by

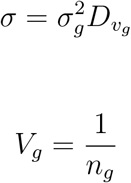

Due to the theoretical predictions of the association between network structure and robustness (Bascompte 2009, Bascompte & Jordano 2007, 2014 and references therein), we also evaluated the correlation between network robustness and nestedness and modularity. To do so, we ran another set of simulations (with 1,000 iterations), across four scenarios, depending on the strength of phylogenetic signal of the species in the higher trophic level (ρ = 1e-04, 1, 2 and 5). Grafen’s ρ varied randomly across the lower trophic level, in each scenario (from 1e-04 to 5). We estimated nestedness using the nested overlap and decreasing fill (NODF) index proposed by Almeida-Neto et al. (2008), and modularity using QuaBiMo algorithm that computes modules, based on a hierarchical representation of species link weights (as we are simulating qualitative networks, all interactions have the same weight) and optimal allocation to modules (Dormann and Strauss 2014).

All numerical simulations and statistical analyses were performed in the R environment (R Core Team 2012) and the simulation code is available at github (https://github.com/bastazini/The-role-of-evolutionary-modes-for-traitdriven-coextinctions-in-mutualistic-networks-network).

## Results

The three scenarios of species extinction were stochastically different, leading to different dynamics of co-extinctions. Networks under a process of random extinctions were more robust than networks experiencing trait-driven cascades (Fig. 2). Despite the smaller differences, networks losing species based on trait distinctiveness were less robust than networks losing species based on size (Fig.2; p(μ_Size trait_ *>* μ_Trait distinctiveness_)= 0.99).

**Fig. 2.**
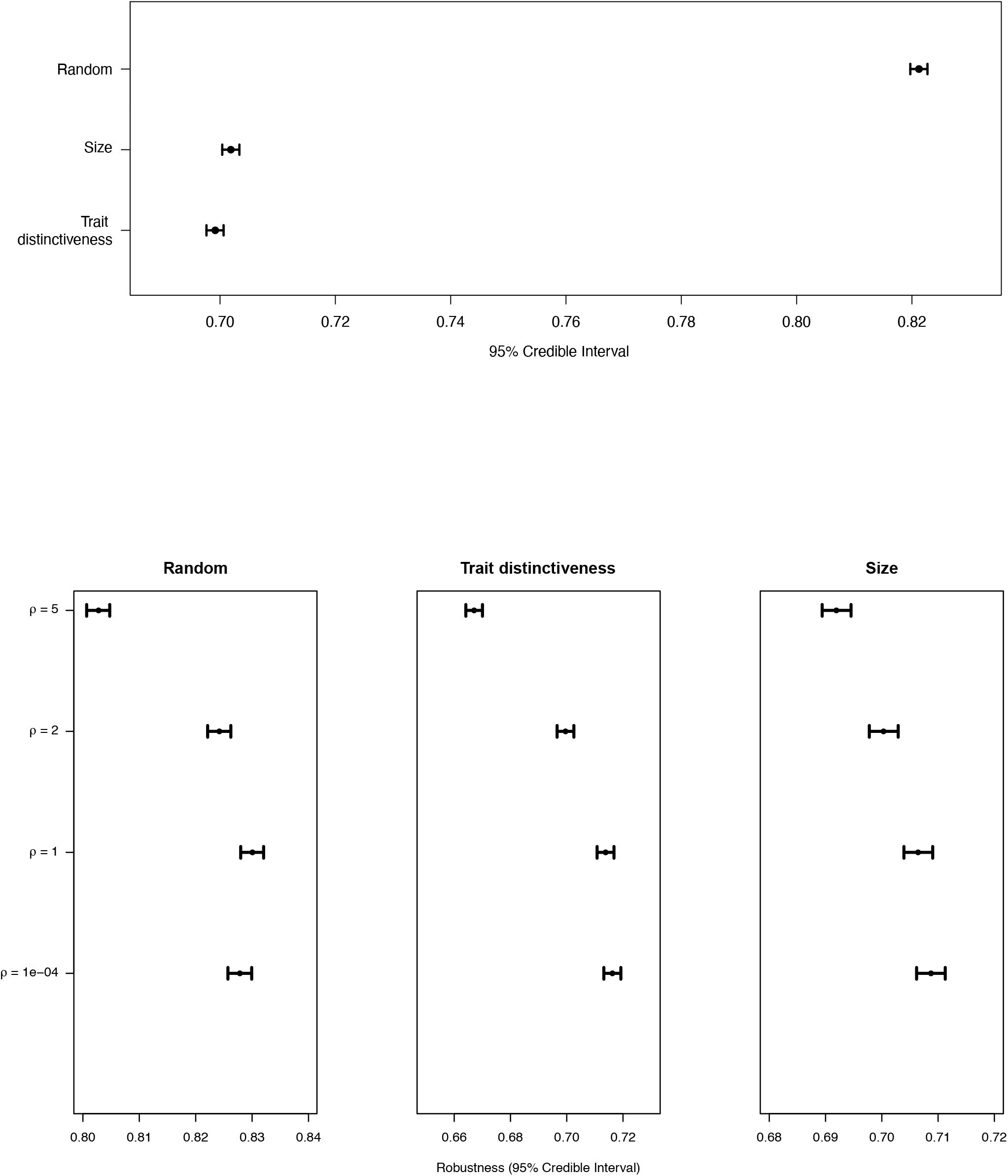
Robustness (95% Credible Interval) for different species elimination schemes, based on trait distinctiveness, trait size and random extinctions.

Our simulations show that the mode of trait evolution matters for network robustness losing functional diversity (Fig. 3). Networks with strong phylogenetic signals were less robust to species extinctions in all three scenarios (Fig. 3). Species traits evolving under Brownian motion led to intermediate levels of robustness in networks undergoing functional attacks (Fig. 3). Our simulations also showed that the phylogenetic signals between interacting species influenced network robustness under functional attacks (Fig. 4). Furthermore, networks losing species with strong phylogenetic signals were less robust to primary species loss in most of the combinations of phylogenetic signals (Fig. 4). The only exception is when the phylogenetic signal in species traits in the lower trophic level is also strong. In this case, there is a large superposition of the posterior distribution in both scenarios of functional attack (Fig. 4).

**Fig. 3.**
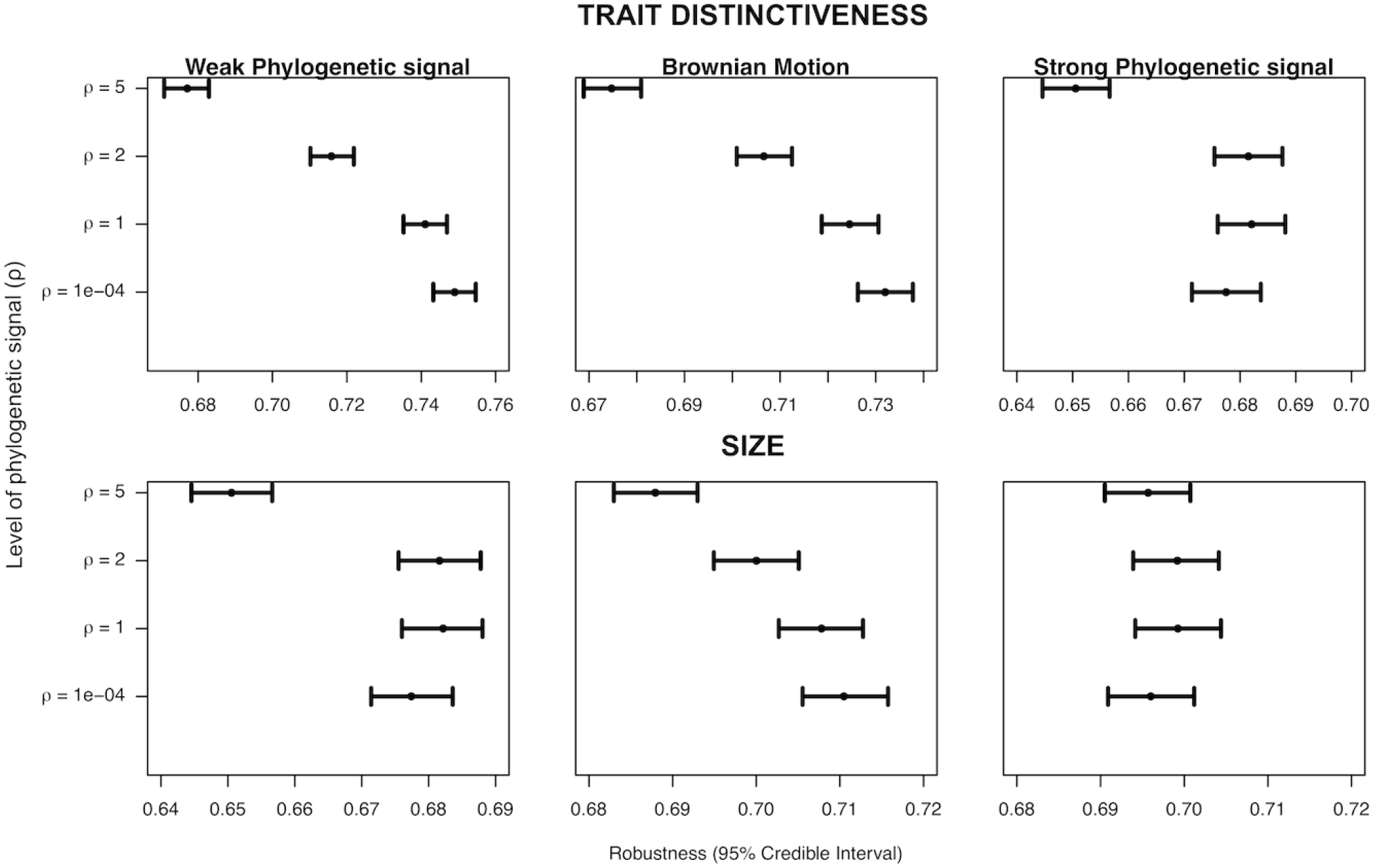
Robustness (95% Credible Interval) for each species elimination schemes, under different phylogentic signal in traits.

**Fig. 4.**
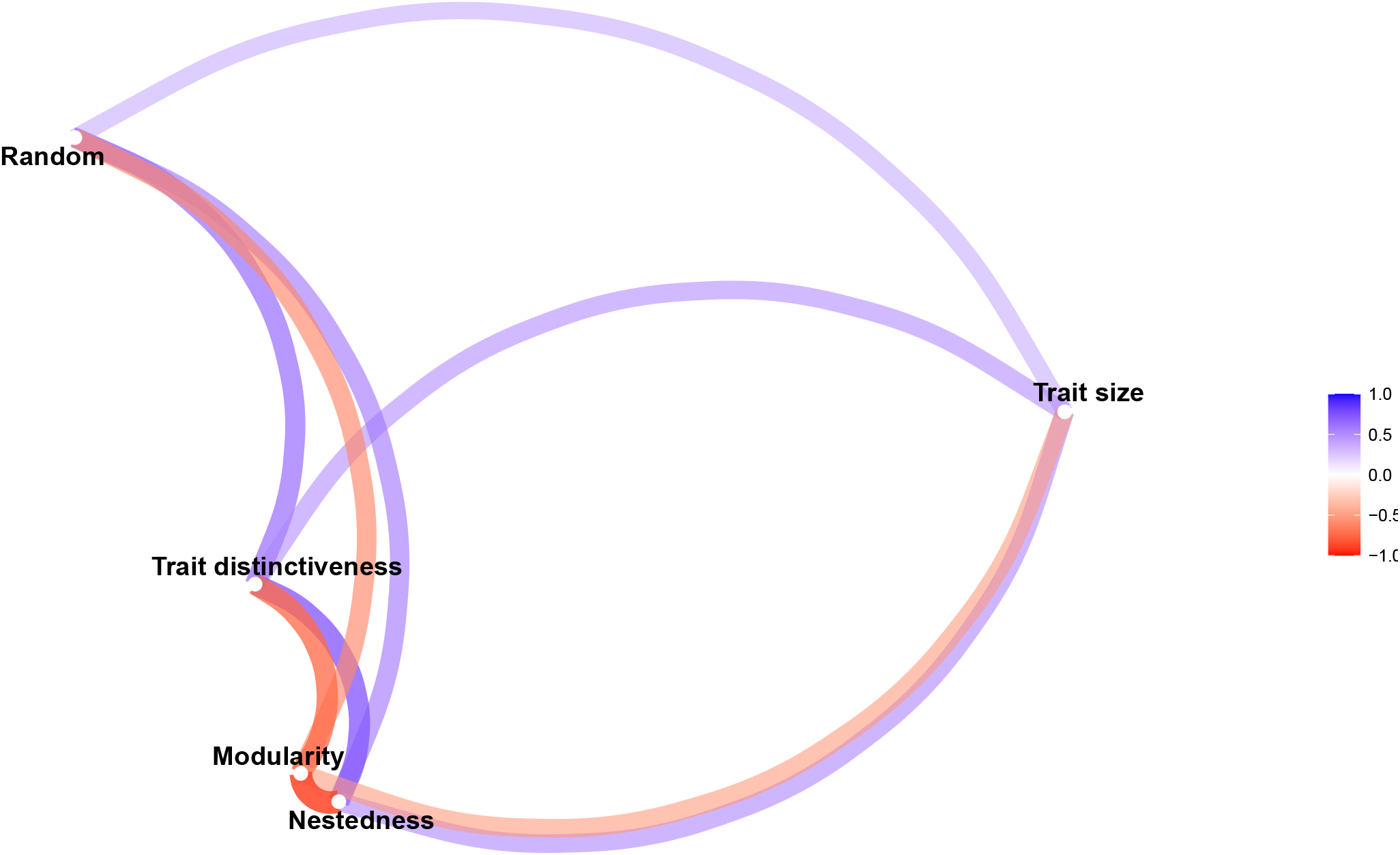
Robustness (95% Credible Interval) for the different evolutionary modes under distinct extinction scenarios in response to each level of phylogenetic signal (ρ) in the other partite (in this case a theoretical lower trophic level).

The association between network robustness and structure behaved similarly across all scenarios, independently of the strength of phylogenetic signal (Fig. 5). Robustness was positively correlated with nestedness (mean correlation ± SD = 0.62 ± 0.10; Fig. 5), and negatively correlated with modularity (−0.63 ± 0.11; Fig 5).

**Fig. 5.** Correlation network, based on Pearson correlation, depicting the association between metrics of network structure (modularity and nestedness) and network robustness across the three scenarios of species loss (Trait size, trait distinctiveness and Random extinctions).

## Discussion

The recent merging of functional and phylogenetic ecology has been contributing to our understanding of the mechanisms underlying species interaction networks (Rezende et al. 2007a,b; Peralta 2016; Bastazini et al. 2017) and the impact of environmental changes on natural communities (e.g., Rezende et al. 2007a; Díaz et al. 2013; Astegiano et al. 2015; Bastazini et al. 2019). Here we evaluated trait-based cascades using minimal model systems of phylogenetically structured mutualistic networks. Although species’ traits can drive organismal and organism-environment interactions in a myriad of complex manners, we explored two specific dimensions of functional diversity expected to have strong consequences to ecological dynamics: body size, a dimension of functional diversity with ubiquitous effect on ecological systems (Purvis et al. 2000; Cardillo et al. 2005; Reynolds et al. 2005; Seguin et al. 2014; Terzopoulou et al., 2015; Verde Arregoitia, 2016; Chichorro et al. 2019) and trait distinctiveness, a facet of functional diversity that warrants unique and/or rare biological interactions and ecosystem functions (Violle et al. 2017). Our results show that extinction cascades based on trait distinctiveness have a more detrimental effect on network robustness, especially when traits evolve under strong phylogenetic signal.

Our results suggest the loss of functional distinctiveness is more detrimental to mutualistic networks than the loss of species with larger size trait values, corroborating empirical results that demonstrate that the loss of more functionally distinct species have a large effect on network robustness (Bastazini et al. 2019). This effect is due to the fact that the role of “functionally unique” species cannot be compensated for by the remaining species in the network (Bastazini et al. 2019; Crooke et al. 2020). Distinct species are irreplaceable components of ecological networks, and yet, still largely ignored in current conservation frameworks (Crooke et al. 2020). Our results support the importance of targeted conservation efforts on species that have unique roles in ecological systems (Crooke et al. 2020).

As phylogenetically related species tend to interact with a similar set of species (Rezende et al. 2007a), we expected that networks exhibiting strong phylogenetic signal would be more robust, as a result of higher trait similarity among species. Contrary to our expectations, this was not the case, and in some situations, strong phylogenetic signal was even associated with reduced robustness. Robustness may be especially reduced when phylogenetic signal in the other set of species is low or when the evolution of traits follows a random walk through evolutionary time in both scenarios of trait based cascades. Scenarios where traits evolve under Brownian motion or traits with weak phylogenetic signal in the lower trophic level suggest that there is a strong coupled phylogenetic response in the set of interacting species, as both scenarios show a proportional response of network robustness with increasing phylogenetic signal. However, when species in the lower trophic level present low phylogenetic signal, network robustness decreases, whereas in the scenario where species in the lower trophic level have strong phylogenetic signal there is not such a clear trend. Rezende et al. (2007a) suggest that ecological communities in which species interactions present a strong phylogenetic component are more likely to suffer co-extinctions following an initial extinction event. Our results corroborate this notion showing that strong phylogenetic signal amplifies the cascading effects of co-extinctions in mutualistic systems.

Although body size has been found to be a fundamental trait capable of predicting species response to environmental gradients (Seguin et al. 2014; Fritschie & Olden 2016) and their extinction risk (Purvis et al. 2000; Cardillo et al. 2005; Reynolds et al. 2005; Terzopoulou et al., 2015; Verde Arregoitia, 2016; Chichorro et al. 2019), its effects depend on the threat and responses and can be fairly inconsistent, as size is represented by different aspects among taxonomic groups (Chichorro et al. 2019). As our simulations are independent of taxonomic identity (and therefore generalist), our finding that trait distinctiveness was more important for robustness further supports the inclusion of other traits, or other facets of functional diversity, rather than size-related traits alone. Therefore the use of size traits as an “all-encompassing trait”, or “key trait” might be misleading in interaction networks or extinction risk studies. Furthermore, empirical evidence from threatened birds and mammals (Crooke et al. 2020) show that species are more ecologically distinct on average which, together with our simulation results, reinforces the need for targeted conservation efforts on species based on their functional distinctiveness (Crooke et al. 2020).

Our simulations support previous findings that robustness should increase with nestedness and decrease with modularity, that network structure can affect its dynamics (Bascompte 2009, Bascompte & Jordano 2007, 2014), and that species phylogenetic relationship affects the degree of ecological network nestedness (Rezende et al. 2007b). Indeed, we found that nestedness had a positive association with the robustness of the system to loss of species or connections. That is likely because a more cohesive structure of nested networks is more redundant, has more alternative states and provides pathways for the persistence of rare species compared to modular ones, and it will not collapse as easily (Bascompte 2009, Bascompte & Jordano 2007, 2014).

We are aware that there are shortcomings to our simulations. First, single trait models may show a poor fit to empirical data (Santamaría & Rodríguez-Gironés 2007; but see Pires et al. 2011). However, trait matching seems to be common in many mutualistic interactions (Garibaldi et al. 2015, Stang et al. 2009, Vizentin-Bugoni et al. 2014, Donoso et al. 2017). Additionally, in our simulations phylogenetic signal is associated with evolutionary process and rate. However, it is important to note that in some situations, this may not be the case, or that this association may be complex (Revell et al. 2008). Other scenarios involving more complex relationships between phylogenetic signal and evolutionary process and rate could bring further insights. Finally, we stress that our framework shares a common shortcoming with similar studies, which assume that a species cannot establish new interactions (“rewire”) in the absence of original mutualistic partners (Vizentin-Bugoni et al. 2020), when secondary extinctions can take place every time a species has no surviving partner (Dunne et al. 2002, Memmot et al. 2004, Burgos et al. 2007, Vieira et al. 2013, Astegiano et al. 2015, Bastazini et al. 2019). Although experimental studies have suggested that rewiring may promote higher resistance in seed dispersal networks (Timoteo et al. 2016, Costa et al. 2018), it should not be common in mutualistic networks with strong trait coupling such as the ones simulated here. That is because trait mismatch prevents new interactions (Santamaría & Rodríguez-Gironés 2007, Bascompte 2009, Vizentin-Bugoni et al. 2014). We still lack a deep understanding of the underlying mechanisms driving rewiring in mutualistic networks. For example, different factors such as spatiotemporal co-occurrence, environmental gradients, and species traits and abundances may determine the probability of species to rewire (Vizentin-Bugoni et al. 2020). The inability to correctly account for the factors determining network rewiring or simulations based on an unconstrained rewiring process could lead to an overestimation of network robustness (Costa et al. 2018), which is undesirable from a conservation point of view.

## Conclusions

Over the past years, ecologists have greatly advanced our understanding of how mutualistic network robustness is associated with phylogenetic patterns (Rezende et al. 2007a; Vieria & AlmeidaNeto 2013; Emer et al. 2019; Bastazini et al. 2019). However, these studies are usually concerned with primary extinctions and the phylogenetic information of just one trophic level or set of species (Vieria & Almeida-Neto 2013; Bastazini et al. 2019). Our simulations demonstrate that cascading effects of co-extinction may spread across taxonomically related species, increasing the erosion of species diversity (Rezende et al. 2007a). Moreover, we show that the interaction between phylogenies of each partite of interacting species may influence network robustness and thus should be considered in studies investigating the association between phylogeny and network robustness. Our models integrating phylogenies of each set of mutualistic species suggest that networks are the most susceptible to collapse when 1) they suffer a targeted attack on more functionally distinct, rather than just larger species, and when 2) species have a strong phylogenetic signal in traits. As a consequence, our results predict that mutualistic networks molded by more recent adaptations are more likely to cope with extinction dynamics than those networks that are based upon conserved traits under trait-based cascades. Despite its simplicity, our *in silico* approach reveals the importance of considering phylogenetic patterns to predict co-extinctions, providing a step forward in understanding cascading effects in natural communities, and developing better conservation strategies.

## Acknowledgments

We thank Rodrigo S. Bergamin, Jeferson Vizentin-Bugoni, André L. Luza and Fernanda Z. Teixeira for kindly reviewing the first draft of this manuscript. We also thank Andreas Kindel, Paulo I. K. L. Prado, Sandra C. Muller for their thoughtful comments and suggestions on the final manuscript. VAGB received support from CAPES (grant #1002302) and is currently funded by National Funds through FCT – Foundation for Science and Technology under the Project UIDB/05183/2020. VPP received support from CNPq, Brazil (grant # 307689/2014-0). PRG was supported by CNPq (307134/2017-2), FAPESP (2018/14809-0), and the Royal Society, London (CHL/R1/180156).

## References

Albert R, Barabási AL (2002) Statistical mechanics of complex networks. Rev Mod Phys 74(1): 47–74

Allesina S, Tang S (2012) Stability criteria for complex ecosystems. Nature 483(7388): 205–208

Almeida-Neto M, Guimarães P, Guimarães Jr PR, Loyola RD, Ulrich W (2008) A consistent metric for nestedness analysis in ecological systems: reconciling concept and measurement. Oikos 117:1227–1239

Astegiano J, Massol F, Vidal MM, Cheptou PO, & Guimarães Jr Pr (2015) The Robustness of Plant-Pollinator Assemblages: Linking Plant Interaction Patterns and Sensitivity to Pollinator Loss. Plos One 10(2): e0117243

Bascompte J (2009) Mutualistic networks. Front Ecol Environ 7: 1–8

Bascompte J, Jordano P (2007) Plant-animal mutualistic networks: the architecture of biodiversity. Annu Rev Ecol Evol Syst 38: 567–593

Bascompte J, Jordano P (2014) Mutualistic networks. Princeton University Press, Princeton

Bastazini VAG, Ferreira PM, Azambuja BO, Casas G, Debastiani VJ, Guimarães Jr Pr, Pillar VD (2017) Untangling the tangled bank: a novel method for partitioning the effects of phylogenies and traits on ecological networks. Evol Biol 44(3): 312–324

Bastazini VAG, Debastiani VJ, Guimarães Jr Pr, Pillar VD (2019).Loss of generalist plant species and functional diversity decreases the robustness of a seed dispersal network. Environ Conserv:1–7.

Brodie JF, Aslan CE, Rogers HS, Redford KH, Maron JL, Bronstein JL, Groves C R (2014) Secondary extinctions of biodiversity. Trends Ecol Evo 29(12): 664–672

Bronstein JF (2001) The exploitation of mutualism. Ecol Let 4:277–287

Burgos E, Ceva H, Perazzo R P, Devoto M, Medan D, Zimmermann M, Delbue AM (2007) Why nestedness in mutualistic networks? J Theor Bio 249(2): 307–313

Burin G, Guimarães Jr Pr, Quental TB (2021) Macroevolutionary stability predicts interaction patterns of species in seed dispersal networks. Science. 372 (6543):733–737

Cardillo M, Mace GM, Jones KE, Bielby J, Bininda-Emonds OR, Sechrest W, Purvis A (2005) Multiple causes of high extinction risk in large mammal species. Science 309(5738): 1239–1241

Ceballos G, Ehrlich PR, Barnosky AD, García A, Pringle RM, Palmer TM (2015) Accelerated modern human–induced species losses: Entering the sixth mass extinction. Sci Adv 1(5): e1400253

Chichorro F, Juslén A, Cardoso P (2019). A review of the relation between species traits and extinction risk. Biol Conserv 273: 220–229

Cohen JE (1977) Food webs and the dimensionality of trophic niche space. P Natl Acad Sci 74: 4533–4536

Cooke, R. S., Eigenbrod, F., & Bates, A. E. (2020). Ecological distinctiveness of birds and mammals at the global scale. Glob Ecol Conserv 22: e00970

Colwell RK, Dunn RR, Harris NC (2012) Coextinction and persistence of dependent species in a changing world. Annu Rev Ecol Evol Syst 43: 183–203

Dethlefsen L, McFall-Ngai M, Relman DA(2007) An ecological and evolutionary perspective on human–microbe mutualism and disease. Nature 449(7164): 811–818

Díaz S, Purvis A, Cornelissen JH, Mace GM, Donoghue MJ, Ewers RM, Pearse W D (2013) Functional traits the phylogeny of function and ecosystem service vulnerability. Ecol Evo 3(9): 2958–2975

Diniz-Filho, AJF, Rangel TF, Santos T, Mauricio Bini L, (2012) Exploring patterns of interspecific variation in quantitative traits using sequential phylogenetic eigenvector regressions. Evolution 66(4): 1079–1090

Donoso I, Schleuning M, García D, Fründ J (2017) Defaunation effects on plant recruitment depend on size matching and size trade-offs in seed-dispersal networks. Proceedings of the Royal Society of London B: Biological Sciences 284: 20162664.

Dormann CF and Strauss R (2014), A method for detecting modules in quantitative bipartite networks. Methods Ecol Evol, 5: 90–98.

Dunn RR, Harris NC, Colwell RK, Koh LP, Sodhi NS (2009) The sixth mass coextinction: are most endangered species parasites and mutualists? P Roy Soc B-Biol Sci 276: 3037–3045

Dunne J, Richard A Williams J, Martinez ND (2002) Network structure and biodiversity loss in food webs: robustness increases with connectance. Ecol Lett 5 (4): 558–567

Eklöf A, Jacob U, Kopp J, Bosch J, Castro-Urgal R, Chacoff NP, Dalsgaard B, Sassi C, Galetti M, Guimaraes Jr PR, Lomascolo SB, Gonzalez AMM, Pizo MA, Rader R, Rodrigo A, Tylianakis JM, Vazquez DP, Allesina S (2013) The dimensionality of ecological networks. Ecol Lett 16: 577–583

Emer C, Galetti M, Pizo MA, Jordano P, Verdú M (2019). Defaunation precipitates the extinction of evolutionarily distinct interactions in the Antropocene. Science advances 5: eaav6699.

Estes, JA, Tinker, MT, Williams, TM, & Doak, DF (1998) Killer whale predation on sea otters linking oceanic and nearshore ecosystems.Science 282: 473–476

Ferriere R, Legendre S (2013) Eco-evolutionary feedbacks adaptive dynamics and evolutionary rescue theory. Philos T R Soc B 368(1610): 20120081

Fritschie, KJ, & Olden, JD (2016) Disentangling the influences of mean body size and size structure on ecosystem functioning: An example of nutrient recycling by a non-native crayfish. Ecology and Evolution, 6(1), 159–169.

Galetti M, Guevara R, Cortes MC, Fadini R, von Matter S, Leite AB, Jordano P (2013). Functional extinction of birds drives rapid evolutionary changes in seed size. Science 340(6136): 1086–1090

Garibaldi LA, Bartomeus I, Bommarco R, Klein AM, Cunningham SA, Aizen MA, Woyciechowski M (2015) Trait matching of flower visitors and crops predicts fruit set better than trait diversity. J Appl Ecol

Grafen A (1989) The phylogenetic regression. Philos T R Soc B 326(1233): 119–157

Guimaraes Jr PR, Sazima C, dos Reis SF, Sazima I (2007) The nested structure of marine cleaning symbiosis: is it like flowers and bees? Bio Lett 3:51–54

Guimaraes Jr PR (2020) The structure of ecological networks across levels of organization Annual Review of Ecology, Evolution, and Systematics 51: 433–460.

Jackson JB, Kirby MX, Berger WH, Bjorndal KA, Botsford LW, Bourque BJ, Warner RR (2001) Historical overfishing and the recent collapse of coastal ecosystems. Science 293(5530): 629–637

Kiers ET, Palmer TM, Ives AR, Bruno JF, Bronstein JL (2010) Mutualisms in a changing world: an evolutionary perspective Ecol Let 13(12) 1459–1474

Lee MD, Wagenmakers EJ (2014) Bayesian cognitive modeling: A practical course. Cambridge University Press, Cambridge

Lefcheck JS, Bastazini VAG, Griffin JN (2015) Choosing and using multiple traits in functional diversity research. Environ Conserv 42: 104–107

May RM (1972) Will a large complex system be stable? Nature 238: 413–414

May RM (2001) Stability and complexity in model ecosystems (Vol. 6). Princeton University Press. Princeton

Memmott J, Waser NM, Price M V (2004) Tolerance of pollination networks to species extinctions. P Roy Soc B-Biol Sci 271(1557): 2605–2611

Minoarivelo HO, Hui C (2016) Trait-mediated interaction leads to structural emergence in mutualistic networks. Evol Ecol 30: 105–121.

Mouquet N, Devictor V, Meynard CN, Munoz F, Bersier L Chave J, Couteron P, Dalecky A, Fontaine C, Gravel D, Hardy OJ, Jabot F, Lavergne S, Leibold M, Mouillot D, Munkemuller T, Pavoine S, Prinzing A, Rodrigues ASL, Rohr RP, Thebault E, Thuiller W (2012) Ecophylogenetics : advances and perspectives. Biol Rev 87: 769–785

Muller-Landau HC, Hardesty BD (2005) Seed dispersal of woody plants in tropical forests: concepts examples and future directions. Cambridge University Press, Cambridge. Book Chapter

Myers JA, Chase JM, Crandall RM, Jimenez I (2015) Disturbance alters beta-diversity but not the relative importance of community assembly mechanisms. J Ecol 103: 1291–1299.

Nee S, May RM, Harvey PH (1994) The reconstructed evolutionary process. Philos T R Soc B 344(1309): 305–311

Neutel AM, Heesterbeek, JA, de Ruite, PC (2002) Stability in real food webs: weak links in long loops. Science 296: 1120–1123

Peralta G (2016) Merging evolutionary history into species interaction networks. Functional Ecology 30: 1917–1925.

Pimm SL (1984) The complexity and stability of ecosystems. Nature307: 321–326

Pimm SL, Jenkins CN, Abell R, Brooks TM, Gittleman JL, Joppa LN, Sexton JO (2014) The biodiversity of species and their rates of extinction distribution and protection. Science 344(6187): 1246752

Pires MM, Koch PL, Farina RA, de Aguiar MAM, dos Reis SF, Guimaraaes Jr PR (2015) Pleistocene megafaunal interaction networks became more vulnerable after human arrival. P Roy Soc B-Biol Sci 282: 20151367

Pires MM, Prado PI, & Guimaraes Jr PR (2011). Do food web models reproduce the structure of mutualistic networks?. PLoS One, 6(11), e27280.

Pocock MJ, Evans DM, Memmott J (2012) The robustness and restoration of a network of ecological networks. Science 335(6071): 973–977

Purvis A, Gittleman JL, Cowlishaw G, Mace G M (2000) Predicting extinction risk in declining species P Roy Soc B-Biol Sci 267(1456): 1947–1952

R Core Team (2012) R: A language and environment for statistical computing R Foundation for Statistical Computing Vienna. URL http://www.R-projectorg/

Redding DW, Hartmann K, Mimoto A, Bokal D, DeVos M, Mooers AO (2008) Evolutionarily distinctive species often capture more phylogenetic diversity than expected. J Theor Bio 251(4): 606–615

Rezende EL, Lavabre JE, Guimaraes Jr PR, Jordano P, Bascompte J (2007a) Non-random coextinctions in phylogenetically structured mutualistic networks. Nature 448(7156): 925–928

Rezende EL, Jordano P, Bascompte J. (2007b). Effects of phenotypic complementarity and phylogeny on the nested structure of mutualistic networks. Oikos 116: 1919–1929.

Revell LJ, Harmon LJ, Collar DC (2008) Phylogenetic signal evolutionary process and rate. Syst Bio 57(4): 591–601

Reynolds JD, Dulvy NK, Goodwin NB, Hutchings J A (2005) Biology of extinction risk in marine fishes. P Roy Soc B-Biol Sci 272(1579): 2337–2344

Santamaria L, Rodriguez-Girones MA (2007) Linkage rules for plant–pollinator networks: trait complementarity or exploitation barriers? PLoS Bio 5(2): e31

Silva W, Guimaraes PR, Guimaraes P, dos Reis SF (2007) Investigating fragility in plant-frugivore networks: a case study for Atlantic Forest. In: Dennis AJ, Schupp EW, Green RJ, Wescott DW. Seed dispersal: theory and its application in a changing world. Wallingford. Wallingford: CAB International. 561–578

Stang M, Klinkhamer PG, Waser NM, Stang I, van der Meijden E (2009) Size-specific interaction patterns and size matching in a plant–pollinator interaction web. Ann Bot 103(9): 1459–1469

Saterberg T, Sellman S, Ebenman B (2013) High frequency of functional extinctions in ecological networks. Nature 499(7459): 468–470

Sazima C, Guimaraes Jr PR, dos Reis SF, Sazima I (2010) What makes a species central in a cleaning mutualism network? Oikos 119(8): 1319–1325

Schleuning M, Frund J, Garcia D (2014) Predicting ecosystem functions from biodiversity and mutualistic networks: an extension of trait-based concepts to plant–animal interactions. Ecography 38(4): 380–392

Schmitz OJ, Buchkowski RW, Burghardt KT, Donihue CM (2015) Functional traits and trait-mediated interactions: connecting community-level interactions with ecosystem functioning. Adv Ecol Res 52: 319–343.

Seguin, A, E Harvey, P Archambault, C Nozais, & Gravel D. (2014). Body size as a predictor of species loss effect on ecosystem functioning. Scientific Reports 4: 4616.

Sole RV, Montoya M (2001) Complexity and fragility in ecological networks. P Roy Soc B-Biol Sci 268(1480): 2039–2045

Strona G & Bradshaw CJ (2018). Co-extinctions annihilate planetary life during extreme environmental change. Sci Rep 8(1): 1–12.

Terzopoulou, S, Rigal, F, Whittaker, R. J, Borges, P A, & Triantis, K A (2015). Drivers of extinction: the case of Azorean beetles. Biol Lett 11: 20150273

Valiente-Banuet A, Aizen MA, Alcantara JM, Arroyo J, Cocucci A, Galetti M, Garcia MB, Garcia D, Gomez JM, Jordano P (2015) Beyond species loss: the extinction of ecological interactions in a changing world. Funct Ecol 29: 299–307

Verde Arregoitia, L.D. (2016), Investigating extinction risk in mammals. Mammal Review 46: 17–29

Vidal MM, Pires MM, & Guimarães Jr, PR (2013) Large vertebrates as the missing components of seed-dispersal networks. Bio Conserv163: 42–48

Vieira MC, Cianciaruso MV, Almeida-Neto M (2013) Plant-Pollinator coextinctions and the loss of plant functional and phylogenetic diversity. Plos One 8(11): e81242

Vieira MC, Almeida-Neto M (2014) A simple stochastic model for complex coextinctions in mutualistic networks: robustness decreases with connectance. Ecol Let 18(2): 144–152

Violle, C, Thuiller, W, Mouquet, N, Munoz, F, Kraft, NJ, Cadotte, M W, … Mouillot, D(2017) Functional rarity: the ecology of outliers. Trends in Ecology & Evolution 32: 356–367

Vizentin-Bugoni J, Maruyama PK, Sazima M (2014) Processes entangling interactions in communities: forbidden links are more important than abundance in a hummingbird–plant network. P Roy Soc B-Biol Sci 281(1780): 20132397

Vizentin-Bugoni J, Debastiani VJ, Bastazini VAG, Maruyama PK, Sperry JH (2020) Including rewiring in the estimation of the robustness of mutualistic networks. Methods in Ecology and Evolution 11: 106–116

Wicksten MK (1998) Behaviour of cleaners and their client fishes at Bonaire Netherlands Antilles. J Nat Hist 32:13–30

